# Tissue-specific consequences of impaired RNA surveillance converge on mitochondrial homeostasis

**DOI:** 10.64898/2026.07.08.737379

**Authors:** Lauryn A. Higginson, Xingjun Wang, Derrick J. Morton

**Author notes:** Mailing Address Corresponding Author: Derrick J. Morton, Molecular and Cellular Biosciences Department of Biological Sciences University of Southern California 1050 Childs Way, RRI 104A, Los Angeles, CA 90089.

## Abstract

RNA surveillance pathways maintain transcriptome integrity by eliminating aberrant, excess, and non-functional RNAs, yet it remains unclear whether distinct tissues exhibit equivalent requirements for RNA quality control. Here, we investigated the tissue-specific consequences of impaired RNA surveillance using a *Drosophila* allelic series of the RNA exosome subunit Rrp40. Comparative transcriptomic analyses revealed that neuronal-enriched head tissue and muscle-enriched thorax tissue exhibit largely distinct molecular programs following reduced RNA exosome activity despite disruption of the same RNA surveillance machinery. Antisense RNAs emerged as particularly sensitive targets of RNA exosome dysfunction, accumulating preferentially in neuronal tissue and largely independent of changes in overlapping sense host transcripts, indicating enhanced requirements for RNA-level quality control within the nervous system. Although tissue-specific transcriptomic alterations diverged substantially, multiple analyses converged on mitochondrial homeostasis as a shared vulnerability. Reduced RNA exosome activity was associated with widespread dysregulation of nuclear-encoded mitochondrial genes, mitochondrial dynamics pathways, and mitochondrial RNA regulatory programs, accompanied by progressive defects in mitochondrial organization, membrane potential, and ATP production. Mitochondrial dysfunction was further associated with activation of proteostatic stress pathways, including p62 accumulation and increased ubiquitination. Together, these findings demonstrate that tissue context shapes the molecular consequences of impaired RNA surveillance while revealing mitochondrial homeostasis as a convergent vulnerability arising from transcriptome instability. More broadly, our findings suggest that distinct tissue-specific defects in RNA regulation converge on common cellular vulnerabilities that ultimately govern tissue homeostasis.

## Introduction

Maintenance of cellular homeostasis requires precise control of RNA abundance and transcriptome composition^1–3^. To maintain transcriptome integrity, RNA surveillance pathways continuously monitor RNAs and eliminate aberrant, excess, and non-functional transcripts that arise during gene expression^4; 5^. This function is particularly important in eukaryotic genomes, where pervasive transcription generates a diverse array of coding and non-coding RNAs, including antisense transcripts, promoter-associated RNAs, and readthrough transcripts (reviewed in^6^). Consequently, RNA decay is not merely a housekeeping process but a fundamental component of gene regulation that shapes transcriptome composition and cellular identity. A key RNA surveillance machinery, the RNA exosome serves as the major 3’-5’ ribonuclease complex responsible for the processing and degradation of a broad spectrum of coding and non-coding RNAs^7^ (reviewed in^8; 9^).

The RNA exosome is a highly conserved multi-subunit complex composed of a catalytically inert core that associates with the ribonucleases DIS3 and EXOSC10 to mediate co-and post-transcriptional RNA processing and decay^10–15^. Together with several cofactors and adaptor complexes^16^, the RNA exosome regulates diverse aspects of RNA metabolism, including rRNA maturation, snoRNA processing, enhancer RNA turnover, and the degradation of cryptic and antisense transcripts^17–20^. Although the molecular composition and biochemical functions of the RNA exosome have been extensively characterized^14; 21^, considerably less is known about how requirements for RNA exosome-mediated surveillance differ across tissues^1; 22; 23^. This question has attracted increasing attention following the discovery that recessive mutations in RNA exosome subunit genes cause a spectrum of tissue-specific disorders with selective effects on a limited number of tissues, frequently including the central nervous system, despite the ubiquitous expression of the RNA exosome (reviewed in^24; 25^). These observations raise a fundamental question: do all tissues depend equally on RNA exosome-mediated RNA surveillance, or do distinct cellular environments exhibit differential requirements for RNA exosome function?

Understanding why some tissues are disproportionately sensitive to impaired RNA surveillance requires identifying the downstream cellular pathway most vulnerable to RNA exosome dysfunction. Mitochondrial homeostasis represents one candidate vulnerability because it requires the coordinated expression of large networks of nuclear-encoded genes^26^. Maintenance of mitochondrial homeostasis requires the coordinated expression of hundreds of nuclear-encoded mitochondrial genes (NEMGs) that regulate mitochondrial biogenesis, metabolism, dynamics, and quality-control pathways^27; 28^. While considerable effort has focused on the transcriptional programs that coordinate nuclear-encoded gene expression^29–32^, comparatively little is known about the post-transcriptional pathways that regulate NEMG expression^29^. Given its central role in co-and post-transcriptional RNA processing and decay, the RNA exosome may contribute to mitochondrial homeostasis^33^ by regulating nuclear-encoded mitochondrial gene expression.

Emerging evidence from model organisms to human cellular systems suggests that the consequences of impaired RNA exosome activity are highly context-dependent^1; 22; 23^. However, most studies have focused on disease-associated phenotypes or individual cell types^22; 23; 34–40^, limiting our understanding of how reduced RNA surveillance impacts distinct tissues within the same organism and whether tissue-specific molecular defects converge on common downstream cellular pathways.

Antisense transcripts represent an attractive class of RNA to investigate tissue-specific RNA surveillance requirements. Antisense RNAs are generated throughout eukaryotic genomes and are frequently targeted for degradation by the RNA exosome and related surveillance pathways^41^. Although many antisense transcripts are rapidly degraded and remain poorly characterized, their accumulation can influence local transcriptional environments and serve as a sensitive indicator of defects in RNA quality-control pathways (reviewed in^42^). Whether impaired RNA surveillance promotes tissue-specific accumulation of antisense transcripts and disrupts mitochondrial homeostasis through altered regulation of nuclear-encoded mitochondrial genes remains unresolved.

Here, we used a *Drosophila* allelic series of the RNA exosome subunit Rrp40 to investigate how tissue context influences the consequences of impaired RNA surveillance. Comparative transcriptomic analyses of neuronal-enriched head and muscle-enriched thorax tissues revealed distinct molecular responses to reduced RNA exosome activity, with antisense transcripts emerging as sensitive targets of RNA surveillance and exhibiting enhanced accumulation in neuronal tissue. Despite substantial tissue-specific divergence in transcriptomic alterations, multiple analyses converged on mitochondrial homeostasis as a common vulnerability, including dysregulation of nuclear-encoded mitochondrial genes and mitochondrial regulatory pathways. Consistent with these findings, *Rrp40* mutant flies exhibited progressive defects in mitochondrial and proteostatic homeostasis. Together, our results demonstrate that tissue-specific consequences of impaired RNA surveillance converge on mitochondrial homeostasis.

## Results

### Reduced RNA exosome activity reveals tissue-specific requirements for RNA surveillance

To investigate whether distinct tissues exhibit differential sensitivity to impaired RNA surveillance, we performed comparative transcriptomic profiling of neuronal-enriched head tissue and muscle-enriched thorax tissue from *Drosophila*, modeling hypomorphic variants in RNA exosome subunit Rrp40^1; 22^. We analyzed two conserved missense substitutions, Rrp40-G11A and Rrp40-G146C (**Fig. 1A**), which were previously shown to reduce Rrp40 abundance *in vivo* and provide an allelic gradient of RNA exosome dysfunction, with G11A representing the more severe and G146C the milder allele^1; 22^. Consistent with these observations, reduced Rrp40 levels are detected in mutant head and thorax tissues relative to wildtype controls (**Fig. S1A-B**). Total RNA isolated from age-matched (Day 1) fly heads and thoraces of wildtype, homozygous G11A/G11A, and homozygous G146C/G146C adult flies was subjected to Nanopore cDNA sequencing to examine tissue-specific transcriptomic consequences of reduced RNA exosome activity (**Fig. 1B**).

**Figure 1.**
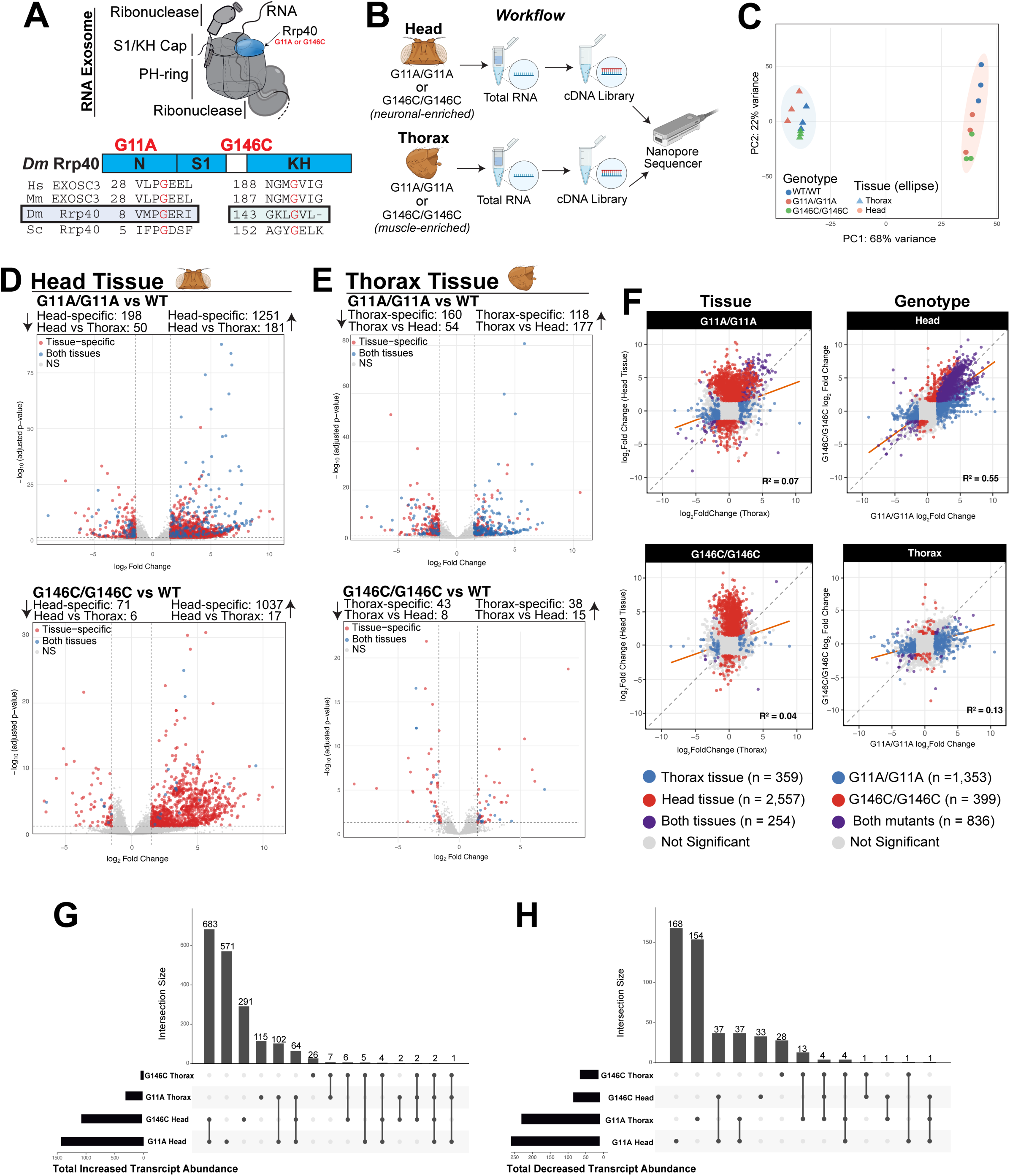
Reduced RNA exosome activity produces tissue-specific transcriptomic alterations. **(A)** Schematic of the RNA exosome complex highlighting cap subunit Rrp40 (light blue) with the location of the disease-associated G11A and G146C missense substitutions below (highlighted in red). Sequence alignment of *Drosophila* Rrp40 and human EXOSC3 demonstrates conservation of variant residues. The G11A substitution affects a highly conserved residue within the N-terminal domain and represents the more severe hypomorphic allele, whereas G146C affects a conserved residue within the KH RNA-binding domain and represents the milder allele, together providing an allelic gradient of RNA exosome dysfunction. **(B)** Experimental workflow. Total RNA was isolated from age-matched (Day 1) adult head (neuronal-enriched) and thorax (muscle-enriched) tissues of *Rrp40* mutant and wildtype flies for Nanopore RNA sequencing. **(C)** Principal component analysis shows that samples are primarily separated by tissue identity, indicating that tissue-specific transcriptional programs dominate transcriptomic variation. **(D)** Differential expression analysis of head (D) tissue comparing *Rrp40* mutants to wildtype controls. Transcripts exhibiting tissue-specific, shared, or non-significant changes are indicated. **(E)** Differential expression analysis of thorax tissue comparing *Rrp40* mutants to wildtype controls. Transcripts exhibiting tissue-specific, shared, or non-significant changes are indicated. **(F)** Correlation of differential expression across tissues and genotypes. Mutant-induced transcriptomic changes are more similar within a tissue than between tissues, highlighting tissue-dependent changes to reduced RNA exosome activity. **(G)** – **(H)** Upset plots showing overlap among significantly increased (G) and decreased (H) transcripts across genotypes and tissues. Most differentially expressed transcripts are unique to a specific tissue-genotype combination.

Principal component analysis shows that samples cluster primarily by tissue identity rather than genotype, indicating that the tissue environment is the dominant determinant of transcriptomic state under conditions of impaired RNA surveillance (**Fig. 1C**). Consistent with this observation, differential expression analysis reveals substantially greater transcriptomic dysregulation in head tissue compared to thorax across both Rrp40 variants (**Fig. 1D-E**). G11A/G11A head samples exhibit the largest number of differentially expressed transcripts, whereas thorax samples show comparatively modest transcriptomic perturbation. These findings suggest that neuronal-enriched tissues possess heightened sensitivity to partial loss of RNA exosome activity.

To determine whether reduced RNA exosome function elicited shared or tissue-specific transcriptomic responses, we compared log_2_ fold change relationships across tissues and genotypes. Comparison of differential expression profiles between head and thorax tissues reveal limited concordance for both Rrp40 variants, suggesting that tissue context is a major determinant of the molecular consequences of impaired RNA surveillance (**Fig. 1F**). In contrast, comparison of G11A/G11A and G146C/G146C within head tissue reveal substantially greater concordance, suggesting that the two alleles perturb overlapping RNA surveillance programs in neuronal tissue despite differences in severity (**Fig. 1F**). Intersection analysis of differentially expressed transcripts further demonstrated that the majority of dysregulated RNAs are tissue-restricted (**Fig. 1G-H**). Upset analysis revealed that transcript overlap was greatest between the G11A/G11A and G146C/G146C alleles within the same tissue, whereas comparatively few transcripts were shared between head and thorax, supporting tissue context as the dominant determinant of RNA exosome-dependent transcript regulation. Together, these findings demonstrate that partial impairment of RNA exosome activity uncovers markedly different tissue-specific requirements for RNA surveillance and RNA decay *in vivo*.

### Antisense transcript accumulation reveals enhanced neuronal requirements for RNA surveillance

To characterize the spectrum of RNAs affected by impaired RNA exosome function, we first classified all significantly increased transcripts by RNA biotype. Protein-coding mRNAs represented the largest class of dysregulated transcripts across all tissues and genotypes, with additional contributions from multiple non-coding RNA classes, including snoRNAs, tRNAs, and lncRNAs (**Fig. S2A-B**). Although antisense RNAs comprised a relatively small fraction of the overall dysregulated transcriptome, their well-established role as direct substrates of RNA exosome-mediated decay prompted us to examine their regulation in greater detail. To further define classes of RNA that are particularly sensitive to reduced RNA exosome activity, we examined antisense transcript regulation across tissues and mutant backgrounds. Antisense RNAs (asRNAs) are established substrates of nuclear RNA surveillance pathways and are frequently targeted by the RNA exosome for degradation^18; 43^. We therefore asked whether impaired RNA exosome function preferentially affects antisense RNA homeostasis *in vivo*.

To conceptualize this framework, we developed a model in which reduced RNA exosome activity leads to the accumulation of asRNAs, resulting in enhanced sensitivity in neuronal tissue (**Fig. 2A**). Examination of representative loci reveals substantial accumulation of antisense transcripts in both *Rrp40* mutant backgrounds, whereas expression of the corresponding sense transcripts remain largely unchanged (Red: Antisense RNA, Blue: Sense Host Transcript) (**Fig. 2B**). Genome browser visualization of representative loci (*FucTc* and *Mpv17*) further confirmed progressive accumulation of antisense transcripts in mutant tissues, with the strongest effects observed in the severe G11A/G11A allele and in neuronal-enriched head tissue (**Fig. 2C**). Together, these data provide direct evidence that impaired RNA exosome function promotes antisense RNA accumulation across multiple gnomic loci.

**Figure 2.**
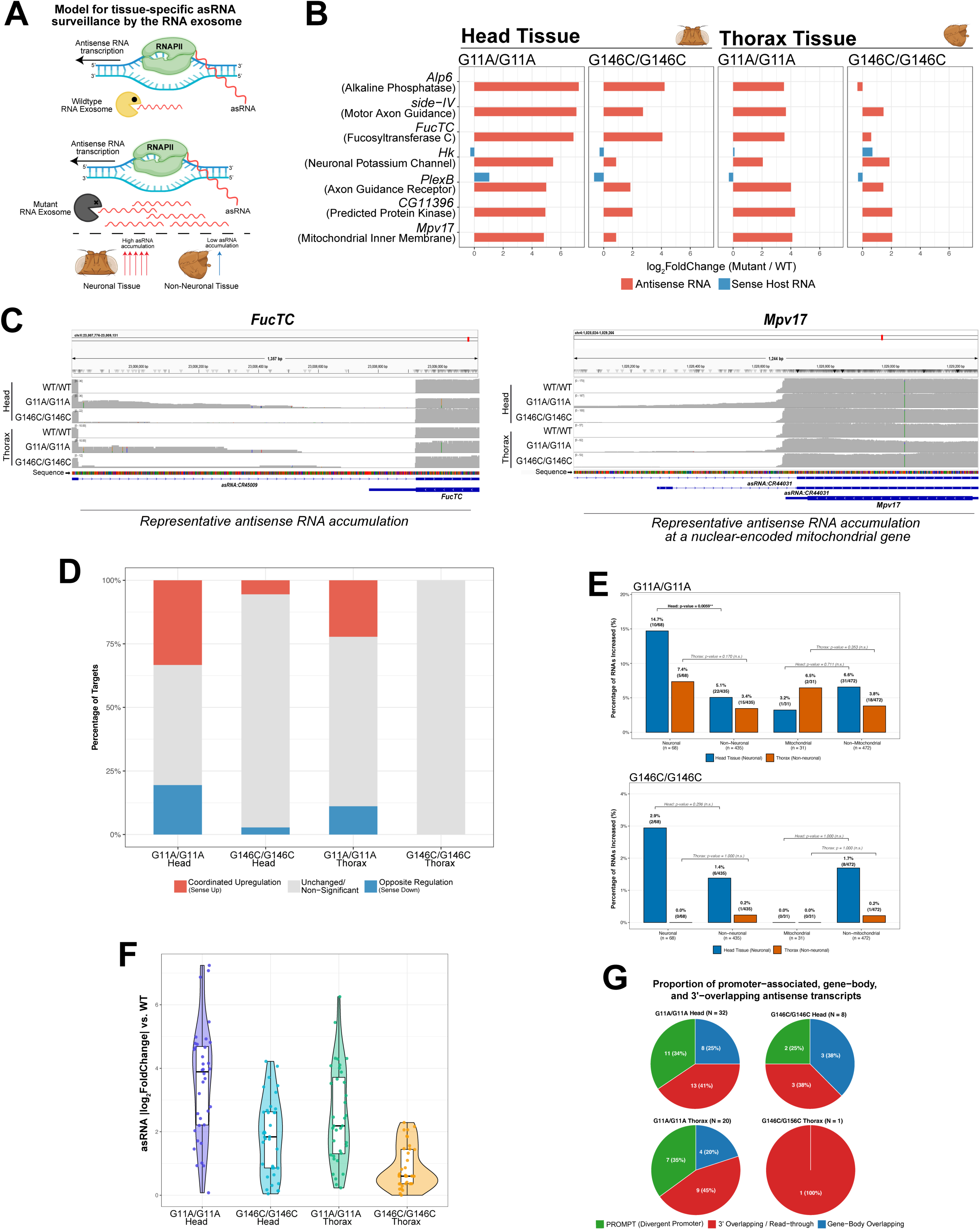
Antisense RNAs are sensitive targets of RNA exosome-mediated surveillance. **(A)** Model illustrating RNA exosome-dependent surveillance of antisense RNAs (asRNAs). Reduced RNA exosome activity results in the accumulation of antisense transcripts, with greater accumulation observed in neuronal than in non-neuronal tissues. **(B)** Representative antisense transcripts exhibiting increased abundance in *Rrp40* mutant head and thorax tissues. Changes in antisense transcripts (red) are shown alongside expression changes of their corresponding sense transcripts (blue). **(C)** Representative genome browser (IGV) views illustrating antisense RNA accumulation at the *FucTC* and *Mpv17* loci. Antisense transcript abundance progressively increases in Rrp40 mutant tissues, with the strongest accumulation observed in the severe G11A/G11A background and in neuronal-enriched head tissue. **(D)** Classification of antisense transcript-host gene relationships. Most increased antisense transcripts accumulate independently of changes in their corresponding sense transcripts, indicating uncoupling of antisense RNA accumulation from host gene expression. **(E)** Distribution of changes in antisense transcript abundance across genotypes and tissues. Antisense RNA exhibits greater accumulation in head tissue relative to thorax tissue, particularly in the severe G11A/G11A mutant background. **(F)** Proportion of increased antisense transcripts associated with neuronal, non-neuronal, mitochondrial, and non-mitochondrial genes. Neuronal-associated antisense transcripts are preferentially increased in head tissue, supporting enhanced requirements for RNA surveillance in the nervous system. **(G)** Genomic classification of increased antisense transcripts. Dysregulated antisense RNAs arise from promoter-associated (divergent promoter), gene-body overlapping, and 3’ overlapping/read-through transcriptional events.

To determine whether antisense RNA accumulation altered the expression of cognate host genes, we compared significantly dysregulated antisense transcripts with their overlapping sense transcripts. Across all tissues and genotypes, most antisense RNAs accumulated without significant changes in host gene expression (**Fig. 2D**). Only a minority of loci exhibit coordinated increases or opposite changes in both antisense and sense transcript abundance, with representative examples shown in **Fig. S2C**. These findings suggest that asRNAs represent particularly sensitive targets of RNA exosome-mediated surveillance and that their accumulation largely occurs independently of host gene transcription.

Consistent with the tissue-specific transcriptomic sensitivity observed in Figure 1, antisense RNA dysregulation is most pronounced in head tissue and in the severe G11A/G11A mutant background (**Fig. 2E**). We next asked whether specific classes of genes were preferentially associated with antisense transcript accumulation. In the G11A/G11A mutant, neuronal-associated loci exhibit a significantly greater frequency of antisense RNA upregulation in head tissue relative to thorax tissue, whereas non-neuronal loci show comparatively modest differences (**Fig. 2F**). These findings suggest that neuronal tissues possess heightened requirements for RNA exosome-mediated antisense RNA surveillance. Notably, *Arc1*, which we previously identified as dysregulated following RNA exosome dysfunction^1; 22^, was again altered in both head and thorax tissues in the present dataset (**Fig. S2D**). This recurrent dysregulation suggests that *Arc1* may serve as a general readout of impaired RNA exosome activity.

Finally, classification of dysregulated antisense transcripts reveals that they originated from diverse genomic contexts, including promoter-associated, gene-body overlapping, and 3’ overlapping/read-through transcripts (**Fig. 2G**). Despite differences in the number of dysregulated antisense RNAs across tissues and genotypes, their genomic distributions remained broadly similar, indicating that impaired RNA surveillance promotes antisense RNA accumulation across multiple transcript classes rather than preferentially a single genomic category.

### Mitochondrial regulatory pathways represent a convergent vulnerability to reduced RNA exosome function

Although transcriptomic consequences to reduced RNA exosome activity differed substantially between tissues, pathway-level analysis revealed a striking convergence on mitochondrial and proteostatic programs. To identify the biological pathways most sensitive to impaired RNA surveillance, we performed gene set enrichment analysis (GSEA) across all tissue and genotype comparisons. Downregulated biological processes are strongly enriched for pathways associated with mitochondrial ATP synthesis, respiratory electron transport chain activity, cytoplasmic translation, and protein folding/refolding (**Fig. 3A**). These pathways are most prominently affected in head tissue, consistent with the enhanced transcriptomic sensitivity of neuronal-enriched tissues to reduced RNA exosome activity. In contrast, upregulated pathways are more variable between tissues and genotypes and include stress-responsive and developmental programs (**Fig. 3A**).

**Figure 3.**
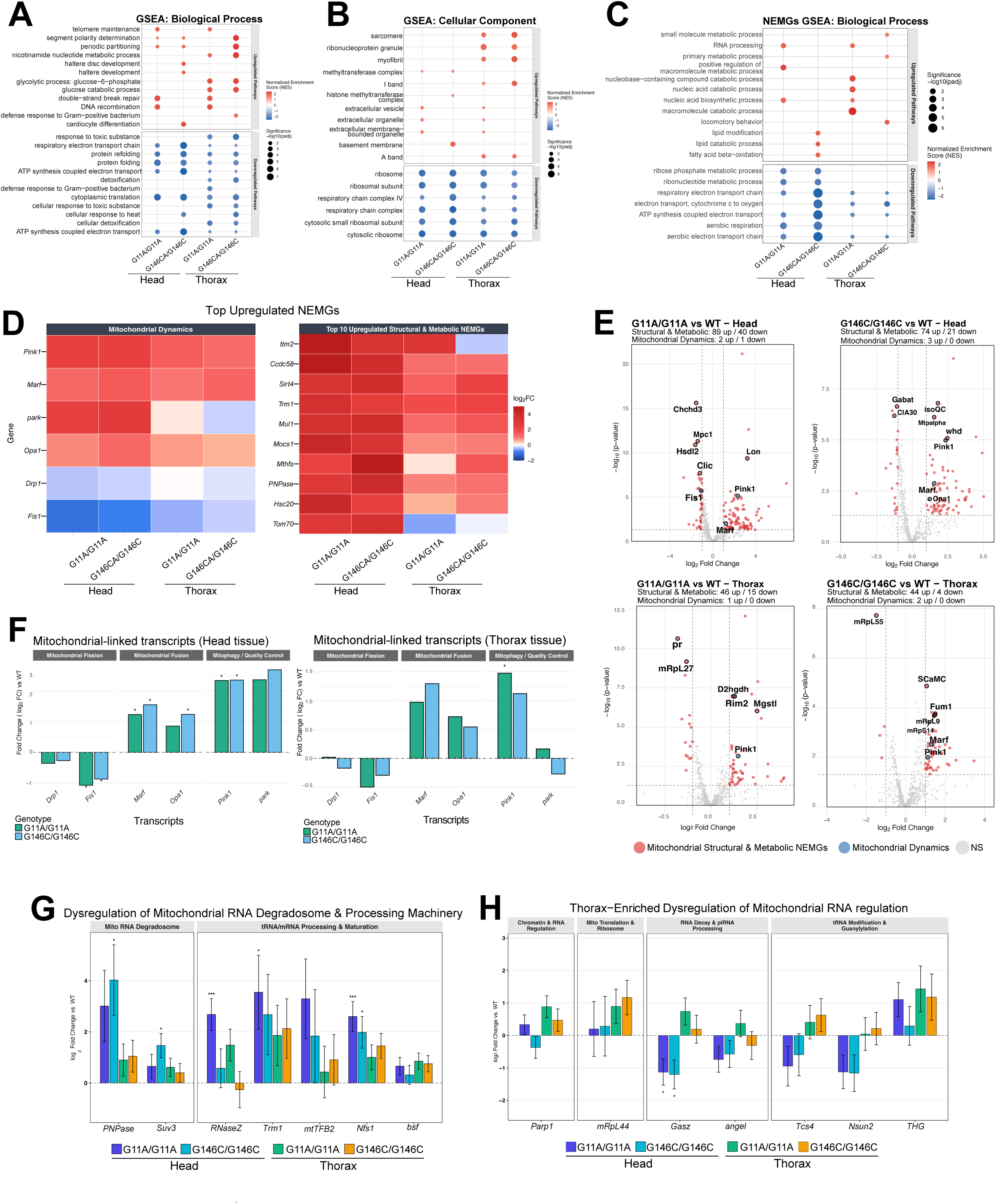
Impaired RNA surveillance remodels mitochondrial gene regulatory networks. **(A-B)** Gene set enrichment analysis (GSEA) of Biological Process (A) and Cellular Component (B) terms associated with transcripts differentially expressed in *Rrp40* mutant head and thorax tissues relative to wildtype controls. Dot size indicates significance (-log_10_ adjusted p-value), and color indicates normalized enrichment score (NES). Mitochondrial and proteostatic pathways emerge as recurrent downregulated processes across tissues and genotypes. **(C)** GSEA restricted to nuclear-encoded mitochondrial genes (NEMGs) identifies depletion of pathways associated with oxidative phosphorylation, ATP synthesis, respiratory electron transport, aerobic respiration, and mitochondrial metabolism in *Rrp40* mutant tissues. **(D)** Heatmaps showing representative dysregulated NEMGs involved in mitochondrial metabolism, respiration, and quality-control pathways across tissues and genotypes. **(E)** Volcano plots highlighting differentially expressed NEMGs in G11A/G11A and G146C/G146C head and thorax tissues relative to wildtype controls. Nuclear-encoded mitochondrial structural and metabolic genes are highlighted with red circles, while mitochondrial dynamics are highlighted with blue circles. Neuronal-enriched head tissue exhibits approximately two-fold more dysregulated NEMGs than muscle-enriched thorax across both *Rrp40* alleles. **(F)** Expression changes in mitochondrial-linked transcripts involved in mitochondrial fission (*Drp1*, *Fis1*), fusion (*Marf*, *Opa1*), and mitophagy/quality control pathways (*Pink*, *park*) in head and thorax tissues. **(G)** Differential expression of genes associated with mitochondrial RNA degradation and processing, including components of the mitochondrial degradosome and mitochondrial RNA surveillance machinery. **(H)** Additional mitochondrial RNA regulatory factors involved in mitochondrial translation, tRNA maturation, and RNA metabolism exhibit tissue-specific dysregulation in *Rrp40* mutant tissues.

Gene ontology cellular component analysis similarly identifies depletion of transcripts associated with respiratory chain complexes, ribosomes, and ribosomal subunits across multiple mutant conditions (**Fig. 3B**). In thorax tissue, enrichment of myofibril-, sarcomere-, and I-band-associated pathways is additionally observed, suggesting that reduced RNA exosome function influences both metabolic and tissue-specific programs. Collectively, these analyses identify mitochondrial homeostasis and proteostatic regulation as recurrent processes vulnerable to impaired RNA surveillance across tissues.

Because mitochondrial pathways emerged repeatedly in enrichment analyses, we next investigated how altered RNA exosome activity affected nuclear-encoded mitochondrial genes (NEMGs). NEMG-specific gene set enrichment analysis reveal significant depletion of pathways associated with aerobic respiration, mitochondrial electron transport, oxidative phosphorylation, and ATP synthesis across both tissues and mutant backgrounds (**Fig. 3C**). Consistent with these pathway-level changes, differential expression analysis identifies widespread dysregulation of NEMGs, including genes involved in mitochondrial metabolism, respiratory chain function, metabolite transport, and mitochondrial quality control (**Fig. 3D-E**). Notably, neuronal-enriched head tissue exhibited approximately two-fold more dysregulated NEMGs than muscle-enriched thorax tissue across both *Rrp40* alleles, further supporting an enhanced requirement for RNA surveillance in neuronal tissues.

To determine whether specific mitochondrial regulatory programs are affected, we examined transcripts involved in mitochondrial dynamics and quality control. Mutant tissues exhibited expression changes consistent with altered mitochondrial remodeling, including reduced abundance of the mitochondrial fission factor *Fis1* together with increased expression of mitochondrial fusion regulators such as *Marf* and *Opa1* (**Fig. 3F**). In parallel, transcripts associated with mitophagy and mitochondrial quality-control pathways, including *Pink1* and *park*, are elevated in mutant animals, particularly in the severe G11A/G11A background.

Given that mitochondrial function depends on extensive post-transcriptional regulation^29^, we next examined genes involved in mitochondrial RNA metabolism. Multiple components of the mitochondrial RNA degradosome and RNA processing machinery, including *PNPase, Suv3*, *RNaseZ*, *mtTFB2*, and *Nfs1*, exhibit altered expression in mutant tissues (**Fig. 3G**). Additional mitochondrial RNA regulatory factors involved in mitochondrial translation, tRNA maturation, and RNA processing are similarly dysregulated (**Fig. 3H**), suggesting broader remodeling of mitochondrial RNA regulatory networks following reduced RNA exosome activity.

Collectively, these analyses demonstrate that, despite substantial tissue-specific divergence in the global transcriptome, impaired RNA surveillance converges on mitochondrial gene regulatory networks. The widespread dysregulation of nuclear-encoded mitochondrial genes, mitochondrial dynamic pathways, and mitochondrial RNA regulatory machinery suggests that reduced RNA exosome activity disrupts mitochondrial homeostasis *in vivo*, thus prompting us to directly examine mitochondrial organization and function in *Rrp40* mutant flight muscle.

### RNA exosome dysfunction disrupts mitochondrial organization, membrane potential, and bioenergetic homeostasis in *Drosophila* flight muscle

To determine whether mitochondrial regulatory signatures identified by transcriptomic analyses were associated with mitochondrial defects *in vivo*, we examined indirect flight muscles (IFMs). IFMs are metabolically active tissues with high energetic demands and are highly sensitive to perturbations in mitochondrial function^44^, as reviewed in ^45^. In *Drosophila*, mitochondrial dysfunction commonly manifests as progressive wing posture defects resulting from impaired flight muscle homeostasis^46; 47^. Consistent with this phenotype, *Rrp40* mutant G11A/G11A flies display progressive wing posture abnormalities, consistent with compromised flight muscle (**Fig. 4A-B**).

**Figure 4.**
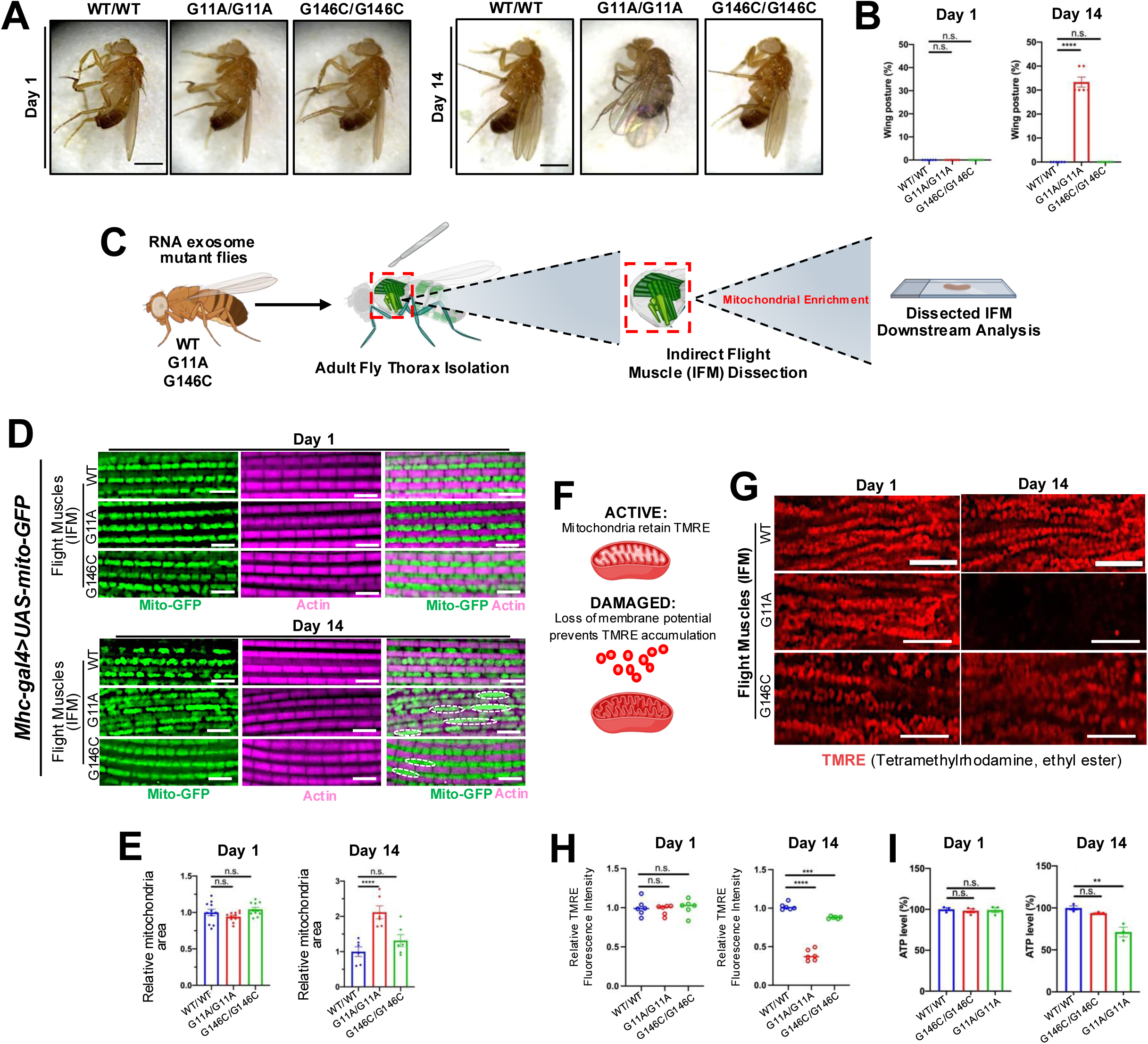
Reduced RNA exosome activity causes progressive mitochondrial dysfunction in vivo. **(A-B)** Age-dependent wing posture in *Rrp40* mutant flies. Representative images (A) and quantification (B) of wing posture abnormalities at Day 1 and Day 14. Defects emerge progressively and are most pronounced in the severe G11A/G11A mutant background. Data are presented as mean ± SD. Statistical significance was determined by one-way ANOVA with Tukey’s multiple comparison test (**p<0.01, ***p<0.001). Scale bars, 10μm. **(C)** Experimental workflow indirect flight muscle (IFM) analysis. Adult thoraces were dissected and IFMs isolated for downstream assessment of mitochondrial morphology and function. **(D-E)** Mitochondrial organization in IFMs was visualized using a mitochondrial GFP reporter and anti-Actin staining. Representative images (D) and quantification of relative mitochondria area (E) at Day 1 and Day 14. White dashed outlines indicate regions or altered mitochondrial morphology, including enlarged, elongated, or disrupted mitochondrial structures consistent with altered fission/fusion dynamics. *Rrp40* mutants exhibit progressive disruption of mitochondrial organization, with the strongest defects in G11A/G11A animals. Data are presented as mean ± SD. Statistical significance was determined by one-way ANOVA with Tukey’s multiple comparison test (**p<0.01, ***p<0.001). Scale bars, 10μm. **(F)** Schematic illustrating assessment of mitochondrial potential using tetramethylrhodamine ethyl ester (TMRE) accumulated in polarized mitochondria, which is reduced following loss of membrane potential. **(G-H)** Representative TMRE images (G) and quantification of TMRE intensity (H) in IFMs at Day 1 and Day 14. *Rrp40* mutants display reduced mitochondrial membrane potential, particularly in the G11A/G11A background. Data are presented as mean ± SD. Statistical significance was determined by one-way ANOVA with Tukey’s multiple comparison test (**p<0.01, ***p<0.001). Scale bars, 10μm. **(H)** (ATP levels were measured from thoracic tissue at Day 1 and Day 14. Reduced RNA exosome activity results in decreased ATP production, consistent with impaired mitochondrial function. Error=SD (**p<0.01, ***p<0.001 One way Anova)

We next examined mitochondrial organization by dissecting IFMs from control and mutant flies expressing a mitochondrial GFP reporter (UAS-mitoGFP, BDSC #8443) driven by *Mhc-Gal4 (*muscle, BDSC #55133). Mitochondria and muscle architecture were visualized using mitoGFP, together with phallodin staining to label F-actin. In wildtype animals, mitochondria are regularly organized between actin-rich myofibrils at both Day 1 and Day 14. In contrast, *Rrp40* mutant muscles display altered mitochondrial organization, with progressive disruption of mitochondrial patterning that became more apparent with age ( **Fig. 4D**). These abnormalities are most pronounced in G11A/G11A animals, which exhibit expanded and disorganized mitochondrial networks by Day 14. In addition, regions highlighted by white dashed outlines showed enlarged, irregular mitochondrial structures that are largely absent in control tissue. These morphological abnormalities are consistent with disrupted mitochondrial dynamics and may reflect an alteration in the balance of mitochondrial fission and fusion. Quantification of mitochondrial area revealed significant changes in mitochondrial architecture in mutant IFMs relative to controls, supporting the conclusion that reduced RNA exosome activity disrupts mitochondrial organization *in vivo* (**Fig. 4D-E**).

To assess whether these structural abnormalities were accompanied by functional mitochondrial defects, we isolated thoraces and enriched mitochondrial fractions from *Rrp40* mutant and control animals for downstream analysis. We then measured mitochondrial membrane potential using tetramethylrhodamine ethyl ester (TMRE), a potentiometric dye that accumulates in polarized mitochondria^48^ (**Fig. 4F**). Compared with wildtype control animals, *Rrp40* mutant samples exhibit reduced TMRE fluorescence intensity, with the most severe loss in G11A/G11A, indicating impaired mitochondrial membrane potential (**Fig. 4G-H**). Notably, membrane potential defects are minimal on Day 1 but become prominent with age, suggesting that mitochondrial dysfunction develops progressively following a reduction in RNA exosome activity.

To determine whether impaired membrane potential is associated with reduced mitochondrial function, we next quantified ATP production in thoracic tissue. Consistent with the TMRE measurements, G11A/G11A mutants exhibit significantly reduced ATP levels at Day 14, whereas ATP production is largely preserved at Day 1 (**Fig. 4I**). These findings indicate that mitochondrial dysfunction extends beyond structural abnormalities and is accompanied by impaired bioenergetic output.

Together, these findings demonstrate that reduced RNA exosome activity disrupts mitochondrial homeostasis *in vivo*. *Rrp40* mutant flight muscles exhibit progressive organismal wing defects, altered mitochondrial organization, and reduced mitochondrial membrane potential, linking impaired RNA surveillance to mitochondrial dysfunction in a high-energy demand tissue. These results provide functional validation of the mitochondrial pathways identified by transcriptomic analyses and establish mitochondrial homeostasis as a key downstream vulnerability of reduced RNA exosome activity.

### Proteostatic stress accompanies mitochondrial dysfunction in *Rrp40* mutant flight muscle

Given the progressive mitochondrial defects observed in *Rrp40* mutant flight muscles, we next asked whether impaired mitochondrial homeostasis is associated with activation of protein quality-control pathways. Mitochondrial dysfunction induces cellular stress responses and can promote the accumulation of damaged or misfolded proteins that require clearance via ubiquitin-dependent degradation pathways and selective autophagy^49–51^. To assess these processes, we examined the accumulation of autophagy adaptor p62/ref(2)P and ubiquitinated proteins within indirect flight muscles (IFMs) of *Rrp40* mutant animals.

We first evaluated p62 accumulation by co-staining IFMs with antibodies against ATP5A to visualize mitochondria and p62. At Day 1, p62-positive puncta are present at low levels across all genotypes (**Fig. 5A-B**). In contrast, at Day 14, G11A mutant muscles exhibit a pronounced increase in p62-positive puncta, whereas WT/WT and G146C/G146C mutant muscles show only sparse labeling (**Fig. 5A-B**). Many p62-positive structures were observed adjacent to or overlapping with mitochondrial networks, consistent with activation of stress-responsive quality-control pathways in tissues exhibiting mitochondrial dysfunction.

**Figure 5.**
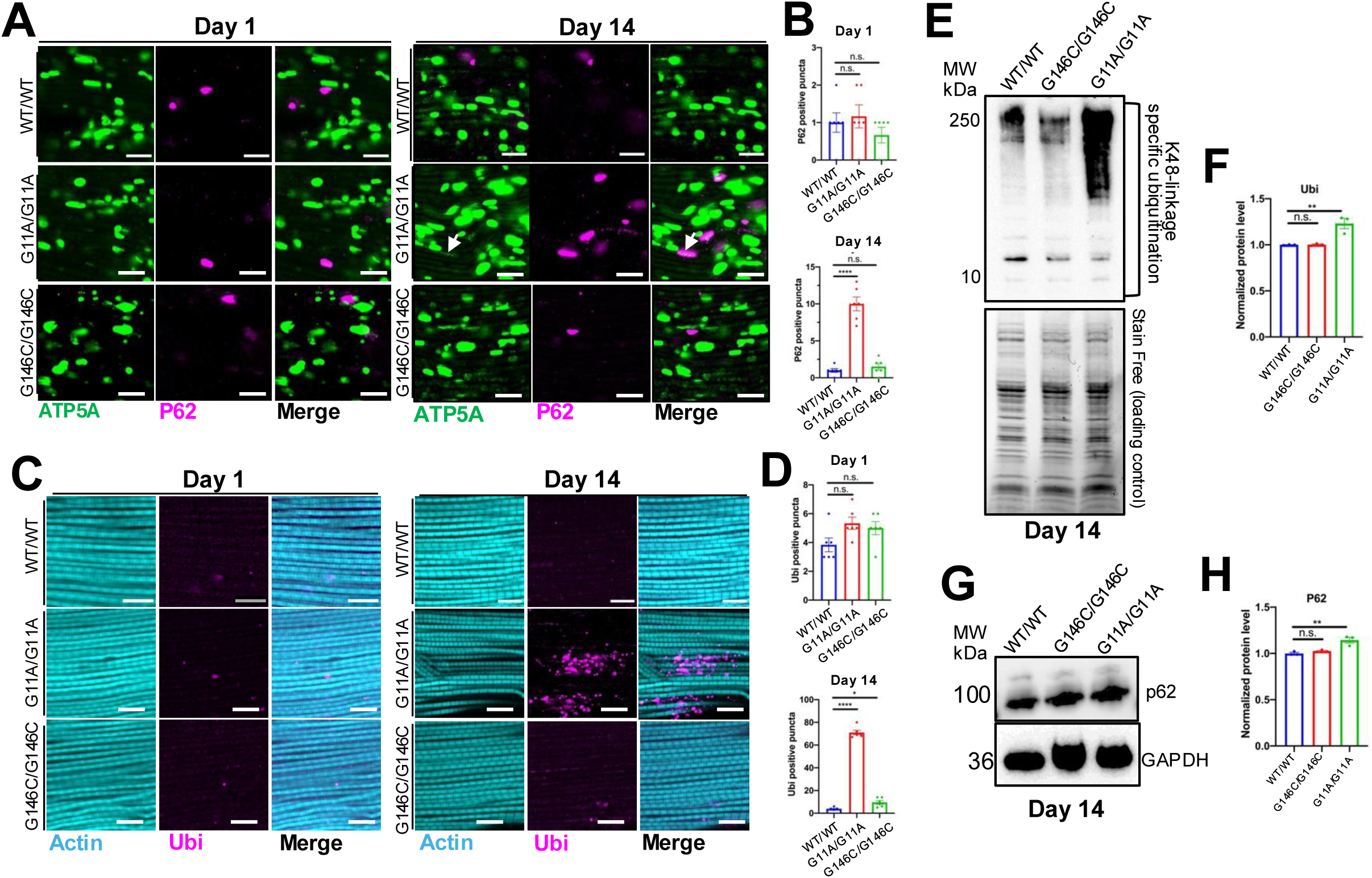
Reduced RNA exosome activity induces proteostatic stress and activation of protein-control pathways. **(A-B)** Accumulation of p62-positive mitochondria in indirect flight muscle (IFMs). (A) Representative confocal images of ATP5A (mitochondria, green) and p62 (magenta) immunostaining IFMs from Day 1 and Day 14 flies. White arrows indicate p62-positive structures associated with mitochondria. *Rrp40* mutant muscles exhibit an age-dependent accumulation of p62-positive mitochondria, with the most pronounced phenotype in the G11A/G11a background. (B) Quantification of p62-positive puncta. Data are presented as mean ± SD. Statistical significance was determined by one-way ANOVA with Tukey’s multiple comparison test (**p<0.01, ***p<0.001). Scale bars, 10μm. **(C-D)** Accumulation of ubiquitinated protein aggregates in IFMs. Representative images of ubiquitin staining (C) and quantification of ubiquitin-positive puncta (D) at Day 1 and Day 14. Increased ubiquitinated protein aggregates are observed in G11A/G11A mutant muscles relative to wildtype and G146C/G146C flies. Data are presented as mean ± SD. Statistical significance was determined by one-way ANOVA with Tukey’s multiple comparison test (**p<0.01, ***p<0.001). Scale bars, 10μm. **(E-F)** Immunoblot analysis of K48-ubiquitinated proteins from thoracic tissue at Day 14. Representative immunoblot (E) and quantification (F) demonstrate increased accumulation of K48-linked ubiquitinated proteins in *Rrp40* mutants, indicative of protein quality control pathways. Data are presented as mean ± SD. Statistical significance was determined by one-way ANOVA with Tukey’s multiple comparison test (**p<0.01, ***p<0.001). **(G-H)** Immunoblot analysis of p62 protein abundance. Representative immunoblot (G) and quantification (H) show the p62 level in *Rrp40* mutant animals relative to the control at Day 14. Data are presented as mean ± SD. Statistical significance was determined by one-way ANOVA with Tukey’s multiple comparison test (**p<0.01, ***p<0.001).

To determine whether these changes were accompanied by broader proteostatic stress, we next examined ubiquitinated proteins in IFMs. Similar to the p62 phenotype, ubiquitin-positive puncta are minimal at Day 1 but accumulated prominently in Day 14 G11A/G11A muscles (**Fig. 5C-D**). Quantification reveals a significant increase in ubiquitinated protein aggregates in G11A/G11A animals relative to both WT/WT controls and G146C/G146C flies, indicating progressive impairment of protein homeostasis following reduction of RNA exosome activity.

To biochemically assess protein degradation pathways, we measured K48-linked ubiquitination, a conserved signal for proteasomal targeting^52^. Immunoblot analysis revealed increased accumulation of K48-linked ubiquitinated proteins in G11A/G11A mutants relative to WT/WT and G146C/G146C animals at Day 14 (**Fig. 5E-F**), consistent with enhanced protein quality control activity and/or impaired clearance of ubiquitinated substrates. We further examined p62 protein abundance and observed a corresponding increase in p62 levels in G11A/G11A mutants at Day 14 (**Fig. 5G-H**), consistent with the age-dependent increase in p62-positive structures observed in IFMs.

Together, these data demonstrate that reduced RNA exosome activity triggers proteostatic stress *in vivo*. Progressive mitochondrial dysfunction in *Rrp40* mutant flight muscle is accompanied by accumulation of p62-positive structures, increased ubiquitinated protein aggregates, and elevated K48-linked ubiquitination, indicating activation of cellular protein quality-control pathways. These findings place proteostasis defects downstream of mitochondrial dysfunction and identify impaired protein homeostasis as an additional consequence of reduced RNA surveillance.

## Discussion

The RNA exosome is a highly conserved RNA surveillance complex that regulates the processing and degradation of diverse classes of RNA^7; 14; 53–57^, yet the extent to which functions of the complex differ across tissues remains poorly understood. In this study, we used a *Drosophila* allelic series of the RNA exosome subunit *Rrp40* to investigate how reduced RNA surveillance impacts distinct tissues. Comparative transcriptomic analyses revealed that neuronal-enriched head tissue and muscle-enriched thorax tissue exhibit markedly different molecular responses to impaired RNA exosome activity despite disruption of the same surveillance machinery. We further identified antisense RNAs as sensitive targets of RNA exosome-mediated decay, with enhanced accumulation in neuronal tissue, suggesting tissue-specific requirements for transcriptome surveillance. Importantly, despite substantial divergence at the level of individual transcripts, multiple independent analyses converged on mitochondrial homeostasis pathways as a shared vulnerability. Reduced RNA exosome activity was associated with widespread dysregulation of nuclear-encoded mitochondrial genes (NEMGs), altered expression of mitochondrial dynamics and quality control factors, and remodeling of mitochondrial RNA regulatory pathways. Consistent with these observations, *Rrp40* mutant animals exhibit progressive mitochondrial dysfunction, characterized by altered mitochondrial organization, reduced membrane potential, and diminished quality-control pathways, including p62 accumulation and increased ubiquitination, indicating a broader disruption of cellular homeostasis. Together, these findings demonstrate that tissue context shapes the consequences of impaired RNA surveillance and reveal mitochondrial and proteostatic dysfunction as common downstream outcomes of reduced RNA exosome activity.

One of the most striking observations from this study is the extent to which tissue context influences the molecular consequences of impaired RNA surveillance. Although the RNA exosome is broadly expressed and performs fundamental functions in RNA metabolism (reviewed in^58^), the transcriptomic consequences of reduced Rrp40 activity differed substantially between head and thorax tissues. Principal component analyses revealed that tissue identity remained the dominant determinant of gene expression, while differential expression analyses demonstrated that individual tissues respond to impaired RNA surveillance through largely distinct transcriptional programs. Similar principles have emerged for other ubiquitously expressed gene regulatory systems, including the spliceosome^59–61^, ribosome^62–64^, and chromatin remodeling complexes^65–67^, where disruption of cellular machines produces unexpectedly tissue-selective phenotypes. Our findings suggest that the consequences of impaired RNA surveillance are not hardwired by the RNA exosome complex, but instead depend on the transcriptomic landscape in which the complex operates. In this framework, the RNA exosome functions with pre-existing tissue-specific gene expression programs, and disruption of RNA surveillance exposes vulnerabilities unique to each cellular environment. While tissue-specific transcriptomic responses to disruption of a ubiquitous RNA surveillance machinery are perhaps not unexpected, the extent of divergence observed between tissues was striking. Most notably, however, these distinct perturbations ultimately converged on a common set of vulnerabilities centered on mitochondrial homeostasis and proteostasis regulation.

Although tissue-specific transcriptomic responses differed considerably, pathway-level analyses repeatedly identified mitochondrial homeostasis as a shared vulnerability. Importantly, this convergence extended beyond pathway enrichment analyses. We observed widespread dysregulation of NEMGs involved in mitochondrial dynamics and quality-control pathways. These findings suggest that impaired RNA surveillance broadly influences the steady-state abundance of nuclear-encoded mitochondrial transcripts and that RNA quality-control pathways contribute to the regulation of mitochondrial gene expression programs. This convergence is notable because the overlap of differentially expressed transcripts between tissues was relatively modest, indicating that distinct molecular perturbations can ultimately impinge upon common cellular processes.

While gene expression is often viewed through the lens of transcriptional regulation^30–32^, our findings underscore the importance of RNA-level quality control in maintaining cellular homeostasis. Antisense transcripts emerged as a particularly sensitive target of the RNA exosome dysfunction and accumulated across tissues and *Rrp40* mutant backgrounds. Importantly, most dysregulated antisense transcripts accumulated independently of changes in their corresponding sense transcripts, suggesting that they are direct or preferential targets of RNA exosome-mediated surveillance rather than secondary consequences of altered transcription. These observations support a growing appreciation that pervasive transcription is not simply tolerated by cells but must be actively managed through dedicated RNA decay pathways. The enhanced accumulation of antisense RNAs in head tissue further suggests that neuronal tissues may place especially stringent demands on RNA quality-control mechanisms. This conclusion is further supported by our NEMG analyses, which revealed two-fold more dysregulated NEMG in neuronal-enriched head tissue than in muscle-enriched thorax tissues in both Rrp40 mutant flies. Given the extraordinary transcriptional complexity of the nervous system, neurons may be particularly dependent on efficient regulation of non-coding and aberrant transcripts to preserve transcriptome integrity.

Mitochondria occupy a central position in cellular physiology, integrating metabolism, signaling, stress responses, and energy production. Maintenance of mitochondrial function requires extensive coordination between nuclear and mitochondrial gene expression programs^29^, making these organelles particularly sensitive to disruptions in RNA metabolism. The observation that mitochondrial RNA processing factors, including components of the mitochondrial degradosome and related RNA regulatory pathways, were also dysregulated suggests broader crosstalk between nuclear RNA surveillance and mitochondrial RNA homeostasis. Growing evidence links defects in RNA processing, RNA transport, and RNA decay to downstream consequences of impaired RNA regulation^68^. Our findings extend these observations by demonstrating that reduced RNA exosome activity disrupts mitochondrial organization, decreases membrane potential, and reduces ATP production *in vivo*. Importantly, these defects emerged progressively with age, suggesting that impaired RNA surveillance compromises the long-term maintenance of mitochondrial function rather than causing an acute developmental defect.

The identification of mitochondrial dysfunction is particularly intriguing given the enhanced sensitivity of neuronal tissue to RNA exosome dysfunction^1; 22; 23^. Mitochondrial abnormalities are a common feature of numerous neurodevelopmental and neurodegenerative disorders and are frequently implicated in the selective vulnerability of post-mitotic tissues^69; 70^. At the same time, muscle and neuronal tissues are among the most energetically demanding cell types in the body, placing extraordinary demands on mitochondrial function and cellular quality-control systems^71; 72^. The disproportionate dysregulation of NEMGs in head tissue suggests that enhanced requirements for RNA surveillance may intersect with heightened demands for mitochondrial homeostasis in neuronal populations. More broadly, the observation that both neuronal and muscle tissues exhibit evidence of mitochondrial dysregulation suggests that mitochondrial dysfunction may serve as a convergent mechanism linking diverse tissue-specific transcriptomic defects to shared cellular outcomes.

Mitochondrial dysfunction rarely occurs in isolation and is frequently accompanied by activation of proteostaic stress pathways^72^. Consistent with this relationship, *Rrp40* mutant muscles accumulated p62-positive structures, exhibited increased ubiquitinated protein aggregates, and displayed elevated levels of K48-linked ubiquitination. These findings indicate activation of protein quality-control pathways and suggest that reduced RNA surveillance ultimately contributes to dysfunction, and that proteostasis has become increasingly apparent in studies of aging and neurodegenerative disease, where defects in one process often exacerbate defects in the other. Impaired mitochondrial function can promote oxidative stress and protein damage, while defective protein turnover can further compromise mitochondrial integrity, creating a self-reinforcing cycle of cellular decline^73^. Our findings suggest that disruption of RNA surveillance may act upstream of these processes by initiating molecular perturbations that ultimately converge on both mitochondrial and proteostaic networks.

Collectively, our findings support a model in which tissue context shapes the consequences of impaired RNA surveillance, while mitochondrial homeostasis emerges as a convergent vulnerability to transcriptome instability. At the same time, our data identify regulatory pathways important for mitochondrial homeostasis - including nuclear-encoded mitochondrial genes, mitochondrial dynamics pathways, and mitochondrial RNA regulatory machinery - as recurrently dysregulated following impaired RNA surveillance. Together, these perturbations converge on mitochondrial dysfunction and proteostatic stress, revealing common downstream consequences of transcriptome instability. These results highlight the importance of RNA quality control pathways in maintaining tissue homeostasis and establish a framework for understanding how disruption of a ubiquitous RNA surveillance machinery can produce distinct molecular consequences yet converge on shared cellular vulnerabilities.

## RESOURCE AVAILABILITY

### Lead contact

- Requests for further information and resources should be directed to and will be fulfilled by the lead contact, Derrick J. Morton.

### Materials availability

- All unique/stable reagents generated in this study are available from the lead contact with a completed materials transfer agreement.

### Data and code availability

- RNA-Seq data have been deposited at NCBI GEO at accession number [GSE337354] and are publicly available as of the date of publication.
- Nanopore-based data is deposited at the NIH Sequence Read Archive (SRA) under BioProject: PRJNA1480847.
- Any additional information required to reanalyze the data reported in this paper is available from the lead contact upon request.

## Acknowledgements

This work was supported by a National Institutes of Health R01 grant (NS131620), Alfred P. Sloan Research Fellowship in Neuroscience award (FG-2023-20698) to D.J.M. L.A.H. was supported by the National Science Foundation Graduate Research Fellowship Program and a University of Southern California Provost Fellowship.

## Author Contributions

Conceptualization, L.A.H., X.W., and D.J.M.; Methodology, L.A.H, X.W., D.J.M, Investigation, L.A.H., X.W.; Writing—original draft, L.A.H. and D.J.M.; Writing—review & editing, L.A.H., D.J.M; Funding acquisition, L.A.H., D.J.M.; Resources, D.J.M; Supervision, D.J.M.

## Material and Methods

### Fly Husbandry and *Drosophila* Lines

All flies were maintained on standard agar/dextrose/cornmeal/yeast medium at 25°C under a 12h light/12 h dark cycle. Unless otherwise indicated, age-matched adult flies were used for all experiments. The Rrp40-G11A/G11A and Rrp40-G146C/G146C lines used in this study were generated previously by CRISPR/Cas9-mediated genome editing (BestGene, Inc., CA) and have been described (ref). These alleles model disease-associated substitutions in the *Drosophila* ortholog of the human RNA exosome subunit EXOSC3. CRISPR-edited control flies were used as controls. To visualize mitochondrial organization in indirect flight muscles, the mitochondrial reporter line UAS-mitoGFP (BDSC# 8443) was introduced into the indicated genetic backgrounds.

### Indirect Flight Muscle Dissection and Immunofluorescence Preparation

Indirect flight muscles (IFMs) were isolated from adult *Drosophila* as previously described^74^ with minor modifications. Adult flies were anesthetized using CO_2_, and the head, wings, legs, and abdomen were removed using forceps while preserving the integrity of the thorax. Isolated thoraces were immediately transferred to 4% paraformaldehyde (PFA) in phosphate-buffered saline (PBS) and fixed overnight at 4°C. Following fixation, thoraces were transferred to PBS containing 0.3% Triton X-100 (PBST). Under a stereomicroscope, the thoracic cuticle was incised along the dorsal midline using fine forceps, and the cuticle was carefully removed to expose the underlying indirect flight muscles. Muscle fibers were gently isolated from surrounding tissue and transferred to microcentrifuge tubes containing PBST using a wide-bore pipette tip generated by trimming the end of a P1000 pipette tip. Isolated muscles were washed three times in PBST for 1 hour each at room temperature to remove residual fixative and prepare samples for subsequent immunofluorescence staining.

### Adult Head Collection for Nanopore Sequencing

Adult flies of the indicated genotypes and ages were anesthetized using CO_2_ and transferred to a chilled dissection surface. Whole heads were separated from bodies using fine forceps under a stereomicroscope. Heads were collected directly into pre-chilled microcentrifuge tubes on dry ice and stored at-80°C until RNA extraction. For each biological replicate, heads were pooled from age-matched flies of the same genotype. Total RNA was subsequently isolated from pooled heads and used for in vitro polyadenylation and Oxford Nanopore cDNA sequencing.

### Oxford Nanopore Sequencing

Adult head (neuronal-enriched) and thorax (muscle-enriched) tissues were dissected from age-matched (Day 1) wildtype, Rrp40-G11AG11A, and Rrp40-G146C/G146C flies. Total RNA was isolated using standard phenol-chloroform extraction methods and quantified prior to library preparation. To enable sequencing of non-polyadenylated transcripts, including antisense RNAs and other RNA exosome-sensitive biotypes, total RNA samples were subjected to *in vitro* polyadenylation using E. coli Poly(A) polymerase (New England Biolabs, M0276S) according to the manufacturer’s instructions. Polyadenylated RNA was subsequently purified by phenol-chloroform extraction followed by ethanol precipitation. Sequencing libraries were generated from approximately 200 ng of in vitro polyadenylated RNA using Oxford Nanopore Technologies (ONT) PCR-cDNA Barcoding Kit (SQK-PCB111.24) following the manufacturer’s protocol. Barcoded libraries were pooled and sequenced on PromethION R10.4.1 flow cells (FLO-PRO114M) using a P2 Solo sequencing device for 24 hours. Raw sequencing data were basecalled in real time using MinKNOW (V6.8.11) with Dorado (v7.11.2) employing default cDNA-PCR basecalling model. Resulting reads were used for downstream transcriptomic analyses, including differential gene expression, antisense transcript identification, gene set enrichment analysis, and characterization of nuclear-encoded mitochondrial gene expression programs.

### Data processing and computational analysis

Reads were aligned to *Drosophila melanogaster* reference genome (FlyBase release r6.66) using minimap2 (v2.30). Gene-level counts were quantified using featureCounts, and genes with fewer than 10 counts across all samples were excluded from downstream analyses. Differential gene expression analysis was performed in R (v4.5.2) using DESeq2 (v1.50.2). The DESeq2 design formula (∼genotype+tissue+genotype:tissue) was used to model the effects of genotype, tissue, and genotype-by-tissue interactions. Gene ontology and pathway enrichment analyses were conducted using clusterProfiler (v4.18.4) with annotations from org.Dme.eg.db (v3.22.0). For antisense RNA (asRNA) analyses, genomic coordinates of annotated antisense transcripts were obtained from FlyBase. Nuclear-encoded mitochondrial genes (NEMGs) were defined using the Gene Ontology term mitochondrion (GO:0005739). Neuronal gene sets were defined using the Gene Ontology terms neurogenesis (GO:0007399) and synapse (GO:0045202).

### Locomotor Function Assay

Negative geotaxis assays were performed as described ^75^ with minor modifications. Newly eclosed *Rrp40* flies (WT/WT, G11A/G11A, G146C/G146C) were collected on day 0, divided into groups of 12, and housed in separate vials. For each trial, age-matched cohorts were transferred into a 25-ml graduated cylinder, and climbing ability was assessed as described^1^.

### TMRE Staining and Quantification of Mitochondrial Membrane Potential

Mitochondrial membrane potential in adult indirect flight muscles (IFMs) was assessed using tetramethylrhodamine ethyl ester (TMRE), a membrane potential-dependent fluorescent dye that accumulates in polarized mitochondria. Adult flies of the indicated genotypes and ages were dissected with TMRE-diluted Schneider’s medium for 20 min at room temperature, protected from light. Following staining, tissues were washed three times in phosphate-buffered saline (PBS) to remove unbound dye and fixed in 4% paraformaldehyde in PBS prior to mounting. Samples were mounted in antifade mounting medium and imaged by confocal microscopy using identical settings. TMRE fluorescence intensity was quantified from IFMs using ImageJ/Fiji. Background fluorescence was subtracted from each image, and mean TMRE intensity was normalized to wildtype controls within each experiment. All image acquisition and quantification were performed using matched settings across samples to permit comparison of mitochondrial potential between genotypes.

### ATP Quantification

ATP levels were measured using the ATP Bioluminescence Assay Kit CLS II (Roche/Sigma-Aldrich) according to the manufacturer’s instructions. Briefly, thoraces from age-matched adult flies of the indicated genotypes were collected and homogenized in ATP extraction buffer. Following centrifugation, supernatants were used for ATP measurements. Bioluminescence was generated by the ATP-dependent luciferase reaction and quantified using a microplate reader. ATP concentrations were determined by comparison to an ATP standard curve generated in parallel for each experiment. Luminance values were normalized to total protein concentration (BCA) and expressed relative to wild-type controls. At least three independent biological replicates were analyzed for each genotype.

### Immunoblotting

Protein lysates (20µg) from fly heads were resolved on 4–20% Criterion TGX polyacrylamide gels (BioRad) and transferred to nitrocellulose membranes. Membranes were blocked for 5-10 minutes in Everyblot Blocking Buffer (BioRad, Cat#12010020) and incubated overnight at 4°C with primary antibodies diluted in blocking buffer. Detection was performed using species-specific horseradish peroxidase (HRP)-conjugated secondary antibodies (Invitrogen) and enhanced chemiluminescence (ECL, Sigma). Primary antibodies include rabbit anti-Rrp40 (custom made with Pacific Immunology), rabbit anti-ATP5A (#ab14748),-GFP (Cell Signaling #2956S), - p62 (#ab178440),-Ubi (#ab140601) and mouse anti-GAPDH (Proteintech).

### Quantification and statistical analysis

Statistical analyses and data presentation were constructed using GRAPHPAD Prism 10 (GraphPad Software, San Diego, CA, USA). All results are represented as mean ± SEM. Statistical significance was determined using either an unpaired t-Test or an ordinary one-way ANOVA. For all statistical analyses, differences were considered significant if *p* < 0.05. Confocal microscopy image processing and quantification were performed using Imaris software (Oxford Instruments), with quantification based on the mean maximum fluorescence intensity per sample.

---

**Supplemental Figure S1.**
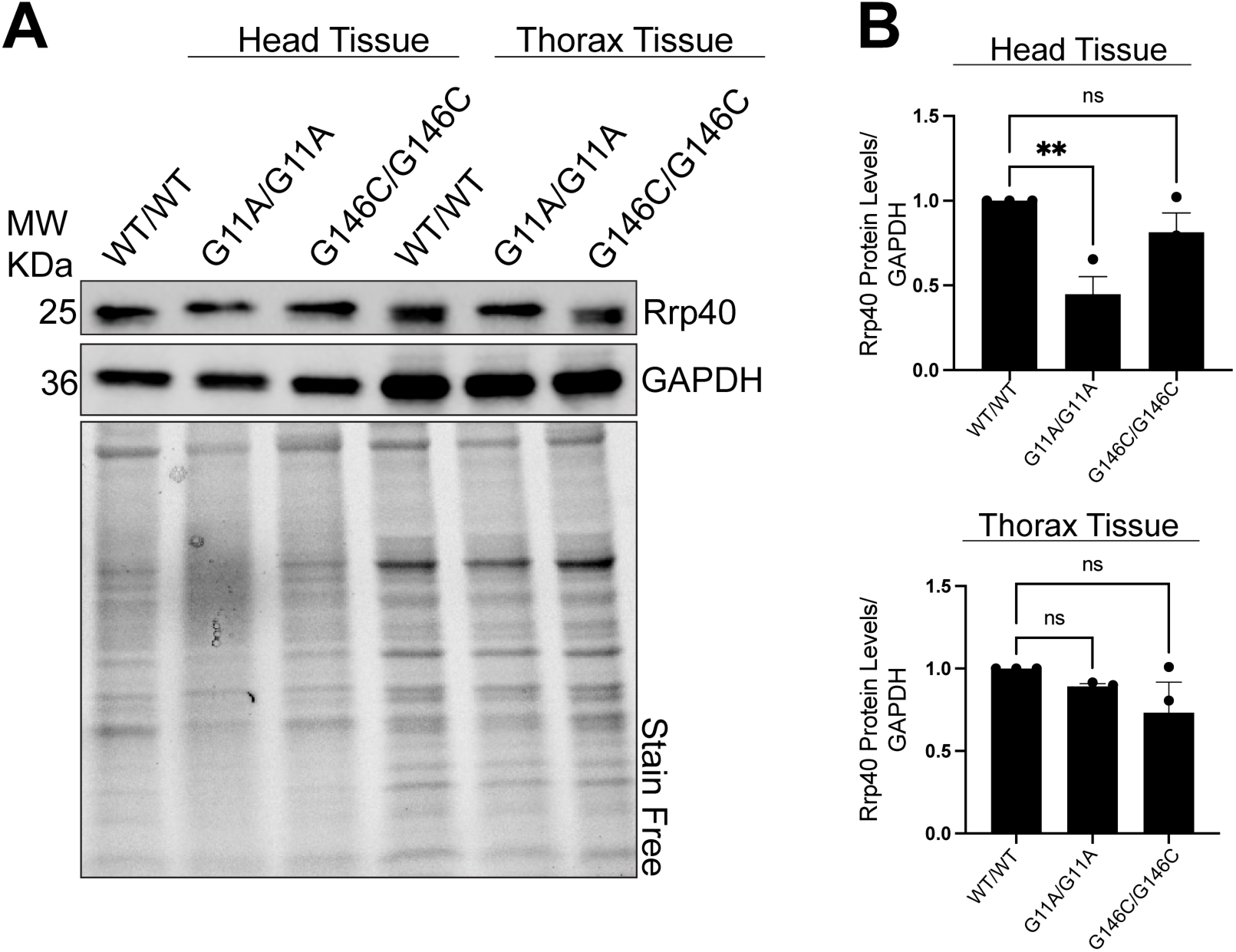
Validation of reduced Rrp40 protein abundance in *Rrp40* mutant tissues. **(A)** Representative immunoblot of Rrp40 protein levels in neuronal-enriched head tissue and muscle-enriched tissue from wildtype (WT/WT), G11A/G11A, and G146C/G146C adult flies. GAPDH served as the loading control. Stain-free total protein imaging is shown as an additional loading control. **(B)** Quantification of Rrp40 protein abundance normalized to GAPDH and compared to WT/WT controls. Data Represent mean ± SEM from n = 3 independent replicates. Statistical significance was determined by one-way ANOVA. *p<0.05; **0<0.01; ns, not significant.

**Supplemental Figure S2.**
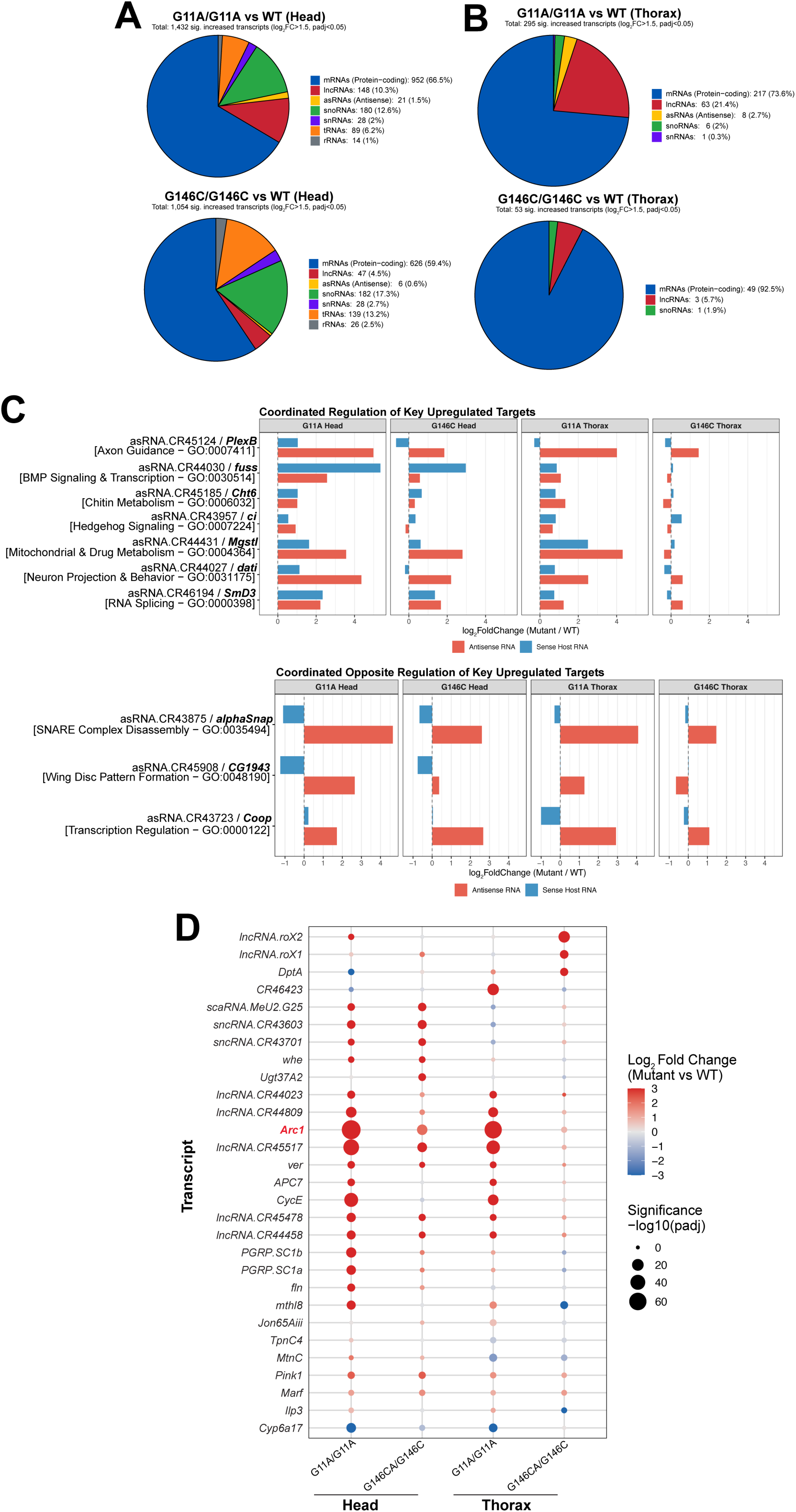
Additional characterization of transcriptomic responses to impaired RNA exosome function. **(A-B)** Classification of significantly increased transcripts by RNA biotype in head (A) and thorax (B) tissues from Rrp40 mutant flies. Protein-coding mRNAs comprise the largest class of dysregulated transcripts, with additional contributions from multiple non-coding RNA classes including lncRNAs, snoRNAs, snRNAs, tRNAs, and antisense RNAs. **(C)** Representative examples of coordinated (top) and opposite (bottom) regulation between antisense RNAs (red) and their corresponding sense host transcripts (blue). Although most antisense RNAs accumulate independently of changes in host transcript abundance (Fig. 2D), a subset of loci exhibit coordinated or opposing changes in steady-state transcript levels. **(D)** Dot plot highlighting representative transcripts recurrently dysregulated following impaired RNA exosome function. *Arc1* (highlighted in red) is reproducibly dysregulated in both head and thorax tissues in the present study and in our previous analyses of *Rrp40*mutant flies^1; 22^. This recurrent dysregulation identifies *Arc1* as a candidate transcriptomic readout of impaired RNA exosome-mediated RNA surveillance.

